# Rapid turnover of corticosterone in humans: the role of a second glucocorticoid hormone

**DOI:** 10.64898/2026.04.27.720824

**Authors:** Mark Nixon, Scott D. Mackenzie, Kerri Devine, Catriona J. Kyle, Rita Upreti, Natalie Z.M. Homer, Rebecca M. Reynolds, Ruth Andrew, Brian R. Walker, Roland H. Stimson

**Author notes:** Corresponding author and person to whom reprint requests should be addressed: Roland H. Stimson.

## Abstract

**BACKGROUND:** Adrenal insufficiency is primarily treated with replacement of cortisol, which is the predominant circulating glucocorticoid. Human adrenals also secrete corticosterone and emerging evidence suggests this may be a safer glucocorticoid replacement therapy. However, little is known about corticosterone in humans, particularly related to its metabolism.

**METHODS:** To investigate the secretion and metabolism of corticosterone in comparison with cortisol, we: 1) investigated the diurnal rhythm of circulating cortisol/ corticosterone in 7 healthy volunteers; 2) quantified A-ring reduction of both hormones in human hepatic cytosol and 3) measured glucocorticoid metabolites in vivo in 24 healthy men; 4) determined the pharmacokinetics of corticosterone via intravenous infusion of 2,2,4,6,6,17α,21,21-[^2^H]_8_-corticosterone; 5) assessed the response of corticosterone and cortisol to 1mcg ACTH in 279 healthy volunteers.

**RESULTS:** The natural diurnal rhythm of corticosterone closely mirrored that of cortisol, and accounted for ∼3% of total circulating glucocorticoid concentrations. Daily corticosterone production, as measured through urinary steroid profiling, was approximately 10-fold lower than cortisol, and corticosterone demonstrated substantially greater metabolism by both 5α- and 5β-reductase than cortisol. In keeping with greater metabolism, the half-life of corticosterone was 28.5 ± 3.3 minutes. Finally, corticosterone demonstrated a greater relative rise in response to ACTH than cortisol, particularly in men, revealing sex-specific differences.

**CONCLUSIONS:** Corticosterone is a dynamic glucocorticoid with faster metabolism and greater response to stimulation than cortisol in humans. These data raise the possibility of distinct roles for these two glucocorticoids and highlight important pharmacokinetic differences with implications for the therapeutic potential of corticosterone replacement in humans.

## Introduction

Glucocorticoids represent a highly effective and widely prescribed class of therapeutics, essential for both suppressing severe inflammation and providing life-sustaining hormone replacement in patients with adrenal insufficiency due to conditions such as congenital adrenal hyperplasia (CAH) ^1,2^. However, long-term exposure to these therapies is linked to severe cardiometabolic toxicities, with an established dose-dependent increase in cardiovascular morbidity and mortality ^3,4^. While recent pharmacological advancements such as sustained-and delayed-release formulations of hydrocortisone (cortisol) have successfully improved the circadian delivery of replacement therapy ^5,6^, these still rely on a glucocorticoid molecule which is associated with cardiometabolic disease. Attempts have also been made to develop selective glucocorticoid receptor modulators in order to separate the beneficial from the toxic effects of glucocorticoids, but this approach has not translated to the clinic to date ^7^. Consequently, achieving optimal disease control without inflicting iatrogenic metabolic harm remains a profound clinical challenge, underscoring the urgent need to identify alternative glucocorticoids with fundamentally safer profiles.

In humans, cortisol serves as the primary glucocorticoid but a second endogenous glucocorticoid, corticosterone, is also synthesised and released from the adrenal cortex. Although corticosterone was investigated in early human glucocorticoid studies ^8^, the relatively low basal plasma concentration (∼5-10% that of cortisol ^9,10^) led to the assumption that corticosterone has a limited, “shadowing”, contribution to glucocorticoid tone, and is often simply considered a mineralocorticoid precursor. However, emerging evidence indicates a stark differentiation in how target tissues handle these two glucocorticoids. Cortisol and corticosterone exhibit distinct susceptibilities to membrane transport by ATP-binding cassette (ABC) transporters, dictating their intracellular availability ^11–17^. ABCB1 (also known as p-glycoprotein), expressed predominantly at barrier sites including the blood-brain barrier, gut and placenta, actively exports intracellular cortisol (and the synthetic glucocorticoids prednisolone and dexamethasone) but not corticosterone ^11,17,18^. Conversely, ABCC1 (also known as multidrug resistance–associated protein 1 (MRP1)), which is more ubiquitously expressed in metabolic tissues including adipose tissue and skeletal muscle, selectively exports intracellular corticosterone but not cortisol ^12,15,17^. This transporter-mediated partitioning provides a theoretical framework for a novel therapeutic: a glucocorticoid capable of penetrating the central nervous system to regulate the hypothalamic-pituitary-adrenal (HPA) axis, while being actively cleared from peripheral tissues susceptible to metabolic toxicity.

A growing body of clinical and preclinical evidence supports this superior therapeutic index. These concepts were initially demonstrated *in vivo* in patients with Addison’s disease, where an acute infusion of corticosterone suppressed ACTH to the same extent as cortisol, while avoiding a comparable induction of glucocorticoid-responsive transcripts in adipose tissue ^15^. Building upon this, a recent placebo-controlled study in patients with CAH demonstrated that an acute infusion of corticosterone suppressed biochemical markers of androgen excess (17α-hydroxyprogesterone, androstenedione, and testosterone), as well as ACTH, similarly to hydrocortisone (cortisol). Crucially, unlike hydrocortisone, corticosterone achieved this without acutely increasing circulating glucose or insulin concentrations ^16^. Furthermore, recent chronic studies in adrenalectomised mice have demonstrated prolonged exposure to exogenous cortisol drives significantly greater insulin resistance compared to equivalent doses of corticosterone ^19^.

Together, these findings suggest that corticosterone replacement may hold benefits over cortisol, particularly in CAH where the balance of HPA axis regulation and side-effects is very difficult to achieve with current therapeutic regimens ^1^. However, the translation of corticosterone into a viable clinical formulation is currently obstructed by a critical knowledge gap. While its acute efficacy has been demonstrated, the fundamental understanding of corticosterone’s basal dynamics in humans and its pharmacokinetic characteristics remains undefined. Establishing these precise parameters is critical for designing appropriate clinical dosing regimens.

Here the dynamic responses of endogenous cortisol and corticosterone to circadian rhythm, and HPA axis suppression and stimulation were studied in humans. Furthermore, key pharmacokinetic parameters including distribution, clearance, and enzymatic metabolism of corticosterone (as a tracer) in humans were compared with cortisol *in vivo* and *in vitro*.

## Materials and Methods

### Diurnal profiling of corticosterone and cortisol

Seven healthy volunteers (4 males/ 3 females) were recruited to a study investigating the diurnal profile of the two circulating glucocorticoids. Inclusion criteria were as follows: aged 18-60 years; no acute or chronic medical condition; not receiving any medication; no glucocorticoid administration by any route within the preceding 3 months; normal screening blood tests (blood count, kidney/ liver/ thyroid function and glucose). Ethical approval was obtained from the ACCORD Medical Research Ethics Committee (AMREC) (16/HV/029) as was consent from each participant. Volunteers attended the Edinburgh Clinical Research Facility (ECRF) after overnight fast in the morning and a cannula sited in the antecubital vein for repeated sampling. Volunteers were given standard meals at 0830, 1200 and 1800 h that were consumed over a 15-minute period (all meals were 55% carbohydrate, 30% fat and 15% protein). Venous blood samples were obtained at 30 minutes intervals from 0800-2200 (a total of 14h), except post-prandially when samples were obtained every 15 minutes for 1 h. Plasma corticosterone and cortisol concentrations were assessed by liquid chromatography tandem mass spectrometry (LC-MS/MS)^16^.

### *In vitro* metabolic clearance

To determine if there was differential metabolism of cortisol and corticosterone, we compared their clearance kinetics *in vitro* using human hepatic cytosol. In addition, the metabolism of D8-corticosterone (as infused previously in human ^15,16^) was compared with that of corticosterone to exclude the possibility of a primary isotope effect as far as possible. The combined metabolism of 5β-reductase and 3α-hydroxysteroid dehydrogenase (3α-HSD) *in vitro* ^20^ was assessed by incubating D8-corticosterone (2,2,4,6,6,17α,21,21-[^2^H]_8_-corticosterone; Cambridge Isotopes), corticosterone, and cortisol with respective radiotracers (1,2,6,7-[^3^H]_4_-corticosterone and 1,2,6,7-[^3^H]_4_-cortisol; Perkin Elmer) with human hepatic cytosol (∼20 mg/mL from adult human donors; Sigma-Aldrich) leading to the formation of 3α,5β-tetrahydrocorticosterone and its isotopically labelled equivalents, as well as 3α,5β-tetrahydrocortisol respectively ^21,22^ with analysis by HPLC with radio-detection. Metabolism of the same steroids by 11β-hydroxysteroid dehydrogenase 2 (11β-HSD2) was assessed in human embryonic kidney cells stably transfected with human *HSD11B2* (HEK-293 11ΒHSD2) as previously described ^23^.

### Assessment of urinary glucocorticoid metabolites in healthy volunteers

To quantify *in vivo* metabolism of corticosterone and cortisol, urine samples from 24 healthy male volunteers were analysed from a subgroup of a previously published study, these 24h urine collections were obtained prior to any intervention ^24^. In brief, inclusion criteria were as follows: aged 20-85 years; BMI <40 Kg/m^2^; no glucocorticoid use in the previous 3 months; no diabetes mellitus or impaired glucose tolerance; no significant hepatic, renal or thyroid disease. Urinary excretion rates of glucocorticoids and metabolites were quantified by GC-MS/MS as described previously ^25^, with the following mass transitions and collision energies (V) used for corticosterone and its metabolites; corticosterone (*m/z* 548→517, 15V), 11-dehydrocorticosterone (*m/z* 474→443, 10V), tetrahydro-11-dehydrocorticosterone (*m/z* 521→400, 15V), 5α/5β-tetrahydro-corticosterone (*m/z* 474→384, 15V), 17-deoxy-β-cortolone (*m/z* 5237→447, 10V), 17-deoxy-β-cortol (*m/z* 531→441, 10V). Glucocorticoid production rates were calculated for corticosterone and cortisol by adding the quantity of the parent hormone and their respective metabolites over the 24h period. The urinary ratios of major A-ring reduced metabolites to their parent hormones (product/ precursor ratios) were also calculated as an *in vivo* index of hepatic 5α- and 5β-reductase activities.

### Pharmacokinetics of corticosterone in healthy volunteers

Three male healthy volunteers were recruited to determine the PK profile of corticosterone. Inclusion criteria were as follows: aged 20-50 years; BMI 20-25 kg/m^2^; on no regular medication and no glucocorticoid administration by any route within the preceding 3 months; normal screening blood tests. Ethical approval was obtained from the South East Scotland Research Ethics Committee 11/SS/0046). Participants attended the ECRF after overnight fast and two cannulae were inserted into antecubital fossa veins, one for intravenous infusion and the other for repeated sampling. D8-corticosterone was administered as a bolus at 0900 h (76.7 µg over 3 minutes from T=0 mins), followed by constant infusion for 3h (7.4 µg/min). The doses infused were designed to achieve circulating D8-corticosterone concentrations of ∼5-10 nM to ensure levels were readily detectable on the LC-MS/MS assay, and were based on our previous human data ^15^. At T+180 minutes, the D8-corticosterone infusion was stopped and blood samples were collected for a further 90 minutes to quantify clearance. Blood samples were collected at 15-minute intervals throughout. Concentrations of corticosterone and D8-corticosterone in plasma were quantified by LC-MS/MS ^15^. Pharmacokinetic parameter were derived according to the following equations:

#### Clearance (*Cl*)

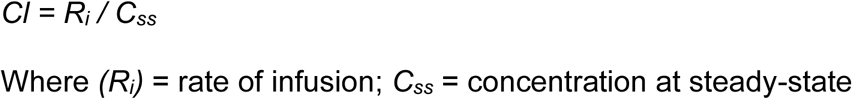

#### Elimination rate constant (*K_el_*)

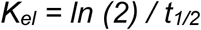

#### First order elimination kinetics

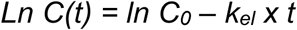

#### Volume of distribution (*V_d_*)

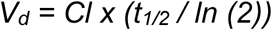

### Response of corticosterone to stimulation of the HPA axis

To validate our pharmacokinetic findings and determine whether corticosterone acts as a highly reactive glucocorticoid *in vivo*, we retrospectively evaluated HPA axis dynamics from a large cohort ^26,27^. Steroid concentrations in plasma samples were revisited from 279 healthy individuals aged 67-78 years (189 males/ 90 females) not prescribed glucocorticoids. At 2200 h the evening before attendance, participants ingested 0.25mg dexamethasone, the following morning a cannula was inserted into an antecubital vein and a baseline blood sample obtained at 0900 h. Thereafter, 1µg of ACTH_1-24_ was administered intravenously, with a blood sample collected 30 minutes later. Pharmacological doses were selected to provide ∼50–75% maximal suppression or stimulation of endogenous cortisol. As plasma cortisol levels had been measured by radioimmunoassay (RIA) and published previously ^27,28^, plasma corticosterone concentrations were also measured by RIA ^22^.

### Data analysis and statistics

Enzyme kinetic parameters were derived using least squares curve fitting (GraphPad Prism V8). The extra sum-of-squares F test was used to compare best-fit parameters between substrates. In the study of HPA activation, concentrations of cortisol and corticosterone were log_2_ transformed and analysed by linear regression with age, gender, BMI and time as co-dependent variables. Student’s *t*-test (paired where appropriate) was used for group comparisons of corticosterone/cortisol ratios and responses to ACTH. Pharmacokinetic analysis was undertaken using Kinetica software (Adept Scientific, Letchworth Garden City, UK). Following discontinuation of D8-corticosterone in steady-state, elimination rate constant (*K_el_*), volume of distribution (*V_d_*) and clearance (*Cl*) were calculated.

## Results

### Plasma corticosterone and cortisol follow a similar diurnal rhythm

In a cohort of healthy volunteers (Table 1), plasma corticosterone concentrations (Figure 1A) mirrored the established diurnal rhythm of cortisol (Figure 1B) over the 14h period, peaking in the morning and reaching a nadir by late evening, with a post-prandial rise in both hormones following lunch ^29^. Consequently, the corticosterone: cortisol ratio remained stable across the study period (Figure 1C). An exploratory sub-analysis by sex revealed no overt dimorphism in these basal rhythms (Supplementary Figure 1).

**Figure 1.**
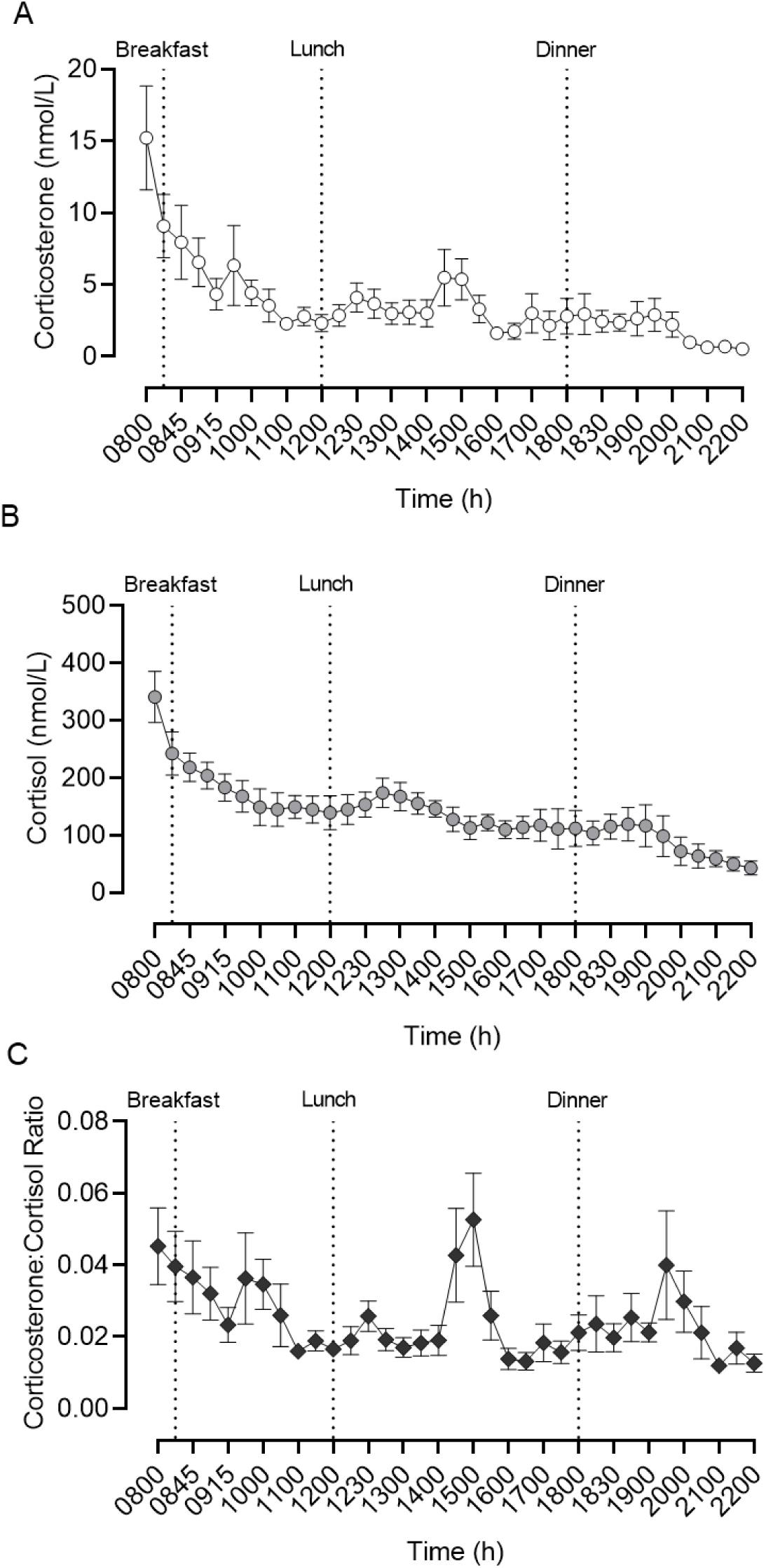
Diurnal profile of glucocorticoids in healthy subjects. A, Plasma corticosterone (open circle). B, Plasma cortisol (grey circle). C, Corticosterone: cortisol ratio in plasma (black diamond). Data are Mean ± SEM. N=7.

**Table 1.**
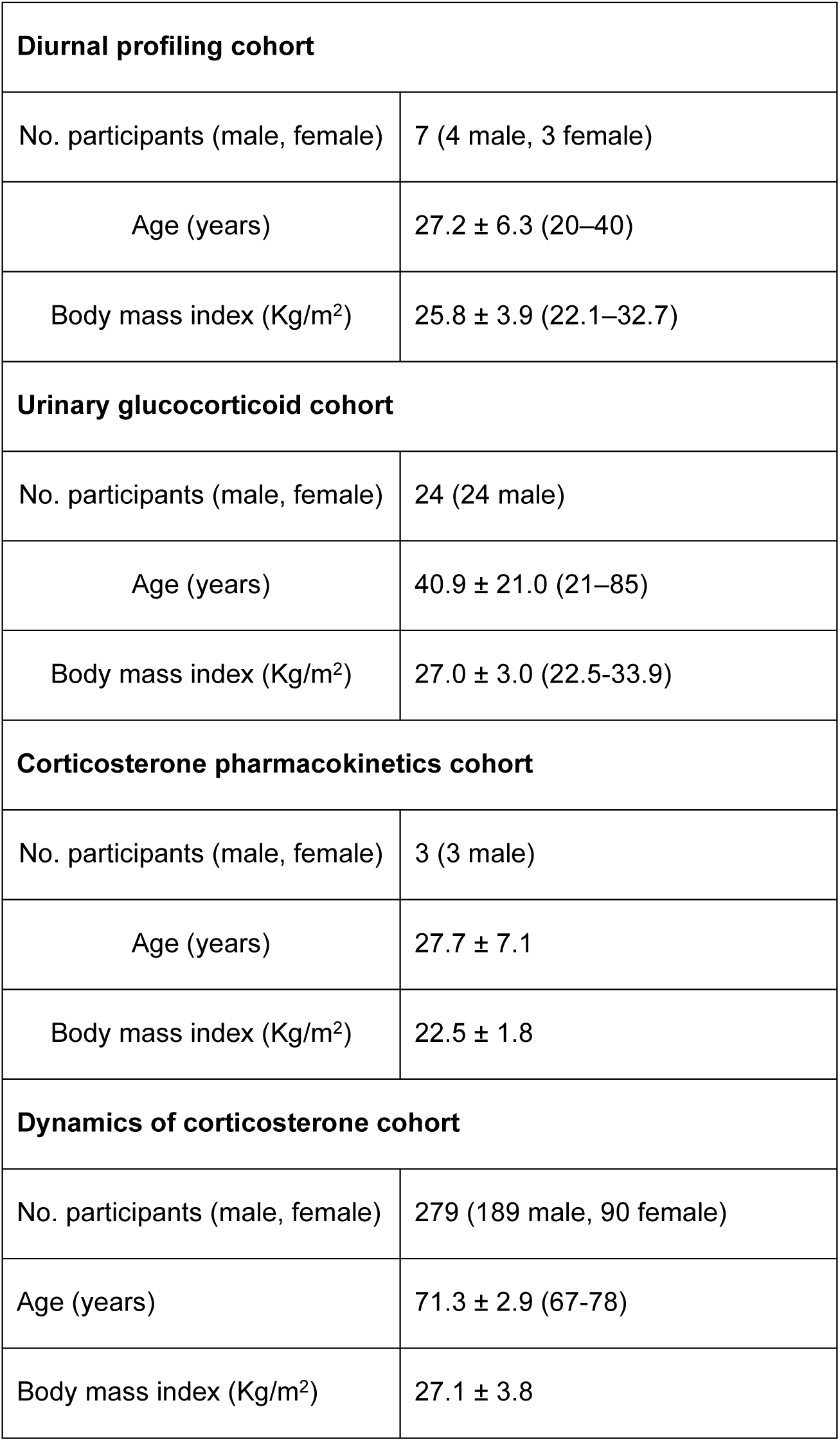
Participant characteristics. *Data are presented as mean ± SD (range) for the 4 separate cohorts studied*.

### Hepatic metabolism of corticosterone is greater than cortisol

To determine if the distinct circulating concentrations of these glucocorticoids are driven by differences in their inherent metabolism, we compared their clearance kinetics *in vitro*. In human hepatic cytosol, reaction velocity of corticosterone and cortisol by 5β-reductase/3α-HSD increased linearly (r^2^ > 0.99) with increasing substrate concentration within the physiological range of both glucocorticoids, indicating non-saturable enzyme kinetics (Figure 2A). In Lineweaver-Burke analysis, the gradient of the regression line (β) for corticosterone (0.0044) was greater than that for cortisol (0.0010) by >4-fold (p<0.001), indicating the rate of metabolism of corticosterone exceeded that of cortisol for a given substrate concentration (Figure 2A). Conversely, for metabolism by 11β-HSD2 in HEK293 cells, metabolic rates were broadly similar for corticosterone and cortisol for a given substrate concentration within the range studied (Figure 2B). Metabolism of corticosterone and D8-corticosterone by 5β-reductase/3α-HSD revealed reaction velocity increased linearly (r^2^ > 0.99) with increasing substrate concentration, and the gradient of the line of best ft for reaction velocity vs. substrate concentration was not significantly different for each substrate (corticosterone: 0.083, D8-corticosterone: 0.080; P=0.13; Supplementary Figure 2A). Both corticosterone and D8-corticosterone were substrates for human 11β-HSD2. No statistically significant differences were found between modelled kinetic parameters for corticosterone (V_max_ = 28.6 ± 2.3 pmol/hr/106 cells; K_m_ = 22.9 ± 3.9 nM) and D8-corticosterone (V_max_ = 24.7 ± 3.6 pmol/hr/106 cells; K_m_ = 17.2 ± 5.5 nM; P = 0.67; Supplementary Figure 2B).

**Figure 2.**
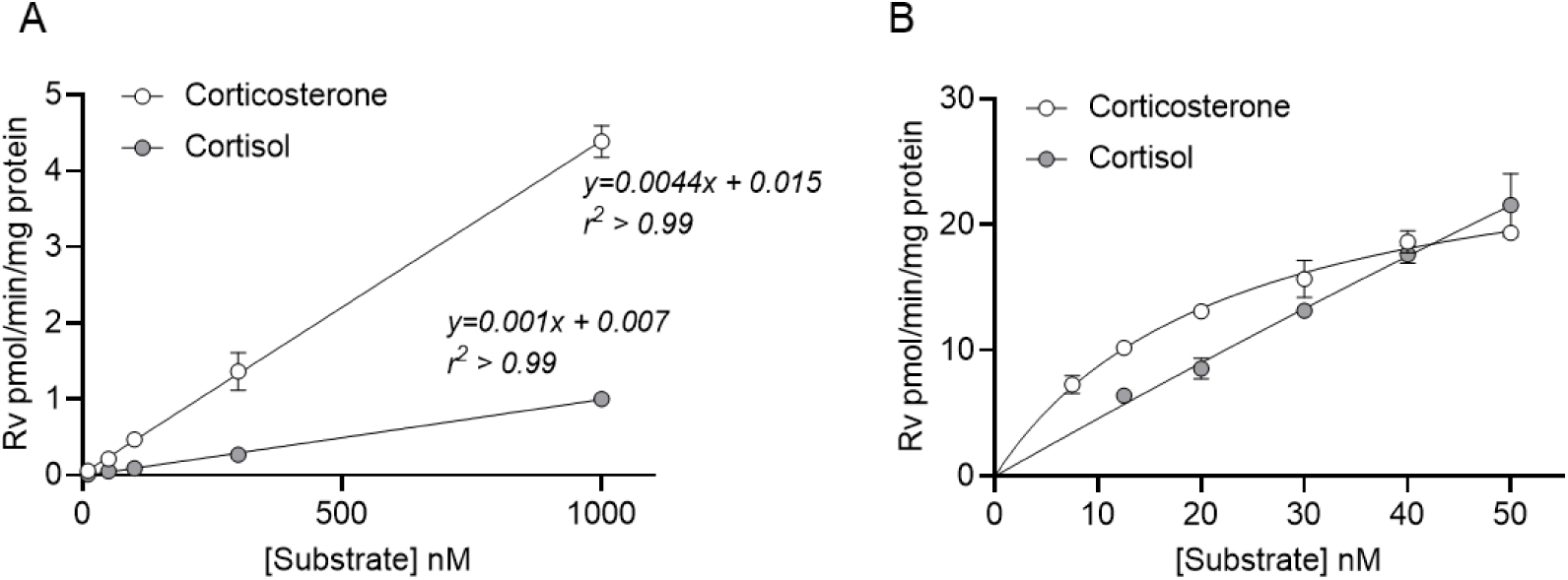
Metabolism of corticosterone is greater than that of cortisol. Reaction velocity (Rv) of: A, Corticosterone (open circles) or cortisol (closed circle s) incubated for 60-240 min with human hepatic cytosol (200 µg protein) and enriched with corresponding [^3^H]_4_-substrate (5 nM). B, HEK293-11β-HSD2 cells incubated (30 min) with 2.5–35 nM corticosterone (open circles) or cortisol (closed circles) and corresponding [^3^H]_4_-substrate (2.5 nM). Data are mean ± SEM. N=3.

### Daily production rate of corticosterone is lower than cortisol and relies on a divergent 5α-reductase clearance pathway

Having established that corticosterone is highly susceptible to 5β-reduction *in vitro*, we next investigated whether this translates to distinct hepatic clearance pathways *in vivo*. In healthy male volunteers (Table 1), the total amount of corticosterone produced, as determined by adding the individual urinary corticosterone metabolites over a 24h-period, was approximately ten-fold lower than that of cortisol (0.65 ± 0.06 vs 7.00 ± 0.43 mg/24 h respectively, P<0.0001; Figure 3A).

**Figure 3.**
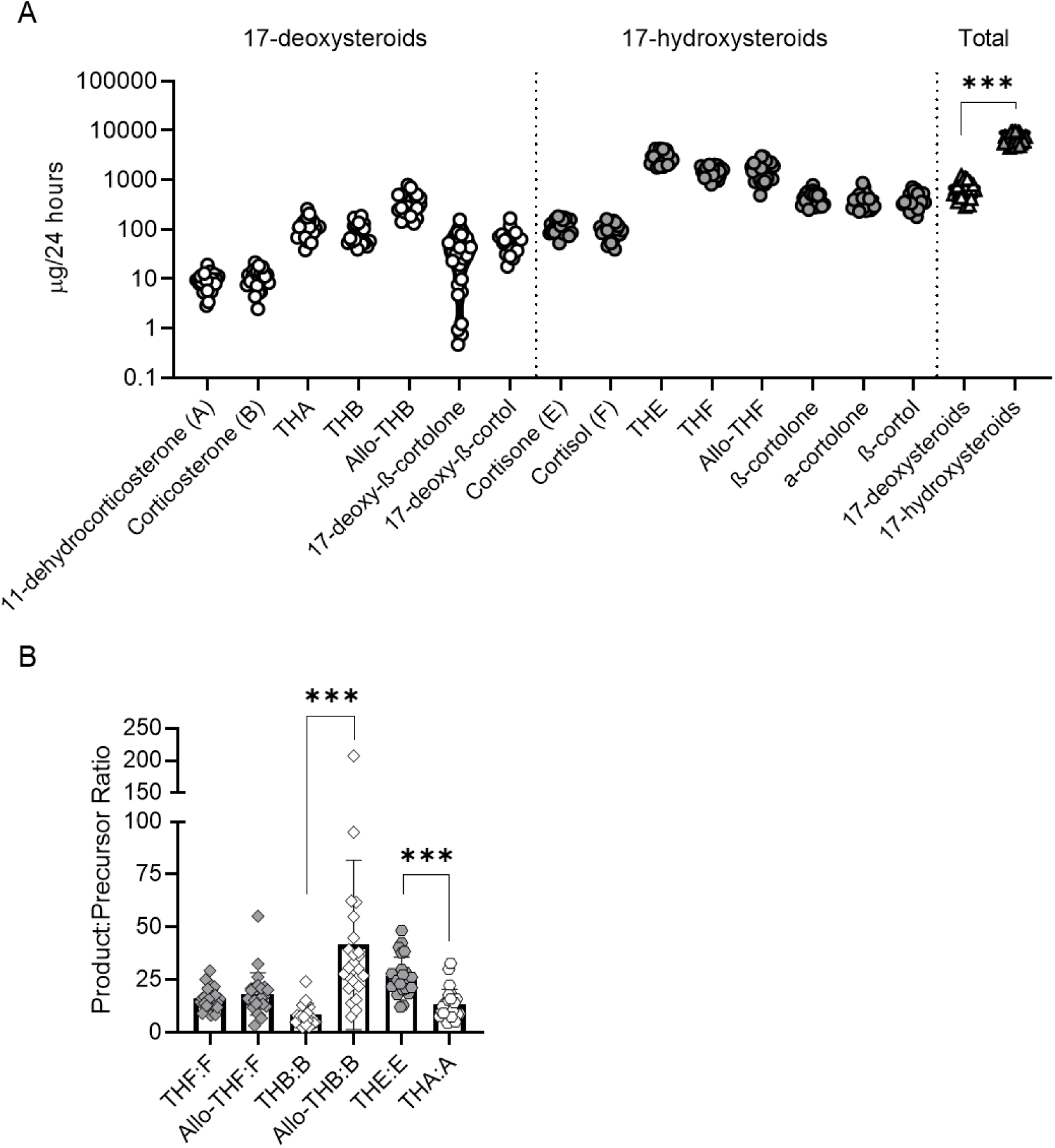
Urinary metabolites of corticosterone and cortisol. Urinary metabolites of corticosterone and cortisol. A, Daily excretion rate of urinary corticosterone metabolites (17-deoxysteroids, white circles) and cortisol metabolites (17-hydroxysteroids, grey circles) from healthy male subjects. Standards were not available for 17-deoxy-α-cortolone, thus for comparison between total 17-deoxysteroids (white triangle) and corresponding total 17-hydroxysteroids (grey triangle), α-cortolone was excluded from the calculation of total daily metabolite excretion “Total”. B, Product: Precursor ratios of A-ring metabolism for corticosterone (white diamonds) and cortisol (grey diamonds). Data are Mean ± SD. N=24. Data for “Total” metabolite excretion and “ratios” analysed by paired t-test. \*\*\**P*<0.001

Beyond production, we utilised this urinary data to investigate the mechanisms driving basal clearance. Cortisol clearance was balanced between the 5α- and 5β-reductase pathways, as evidenced by similar ratios of allo-tetrahydrocortisol (allo-THF) to cortisol compared to THF to cortisol (18.3 ± 1.7 vs. 16.3 ± 1.3 respectively; P=0.2175; Figure 3B). In contrast, the 5α-reduced metabolite of corticosterone, allo-tetrahydrocorticosterone (allo-THB), constituted the vast majority of corticosterone A-ring reduction, resulting in a significantly higher allo-THB/B ratio compared to the THB/B ratio (41.6 ± 6.4 vs. 8.6 ± 1.1, respectively; P<0.0001). We also evaluated 5β-reduction of their respective 11-keto metabolites, 11-dehydrocorticosterone (A) and cortisone (E), which are restricted from utilising the 5α pathway. Indeed, the *in vivo* 5β-reduction of 11-dehydrocorticosterone was half that of cortisone (THA/A: 13.4 ± 0.9 vs THE/E: 26.7 ± 1.9; P<0.0001). These data show that corticosterone is preferentially metabolised by 5α-reductase rather than 5β-reductase *in vivo*.

### Corticosterone is rapidly cleared *in vivo*

Having identified a highly active, 5α-reductase-driven clearance pathway for corticosterone *in vivo*, we next sought to definitively quantify the pharmacokinetic consequences of this divergent metabolism. To achieve this, *in vivo* stable-isotope dilution at steady-state was performed using a continuous intravenous infusion of D8-corticosterone following an initial IV bolus in three healthy male subjects (Table 1). D8-corticosterone concentrations reached steady state by t=∼105 minutes, with a mean concentration of 24.5 ± 3.7 nmol/L between t=105-180 mins (Figure 4A).

**Figure 4.**
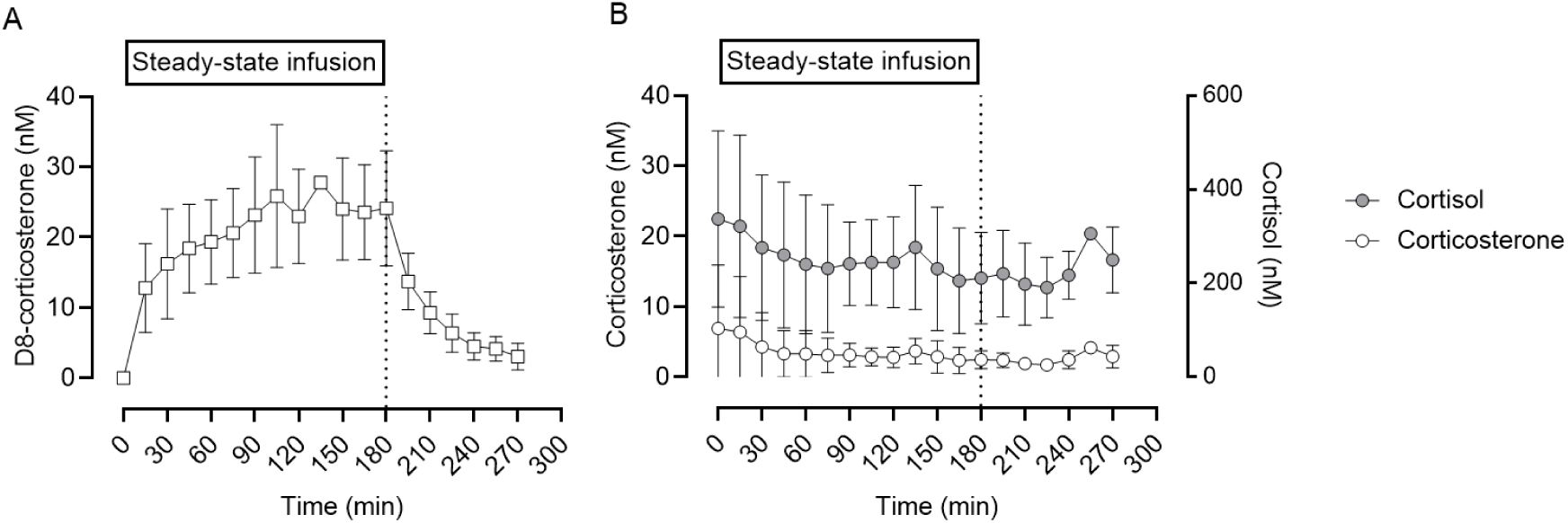
Pharmacokinetic profiles of intravenous D8-corticosterone *in vivo*. Steady state infusion for 180 mins (T0 = 0900 h); N=3. A, D8-Corticosterone (open squares). B, endogenous corticosterone (open circles); endogenous cortisol (closed circles). Data are mean ± SD.

Following discontinuation of the D8-corticosterone infusion, the decay was modelled using first-order kinetics, with an elimination rate constant (K_el_) of 0.031 ± 0.014 min^-1^ and t_½_ 28.5 ± 3.3 mins. The steady state clearance was calculated to be 0.96 ± 0.20 L/min. Finally, the apparent volume of distribution (V_d_)_,_ derived from clearance and t_½_ using the steady state parameters, was 39.6 ± 5.3 L.

### Plasma corticosterone is more sensitive to HPA axis activation than cortisol

Given the rapid *in vivo* clearance of corticosterone established by our stable-isotope pharmacokinetic data, we hypothesised that it would act as a highly dynamic "fast-responder" during acute physiological stress. To investigate this, we evaluated the adrenal regulation of both glucocorticoids in a large cohort of subjects who underwent dynamic testing of the HPA axis through dexamethasone suppression followed by low-dose ACTH stimulation (Table 1) ^27,28^. Both plasma corticosterone and cortisol concentrations increased significantly following ACTH_1-24_ stimulation (Figure 5A,C). However, reflecting its high-turnover kinetics, corticosterone exhibited a disproportionately greater stress response compared to cortisol, nearly double that of cortisol (4.2 vs 2.4-fold increase, respectively; *P*<0.001). Consequently, the circulating corticosterone:cortisol ratio increased significantly following HPA axis activation, demonstrating a shift in the systemic balance of glucocorticoids during acute stress (Figure 5E).

**Figure 5.**
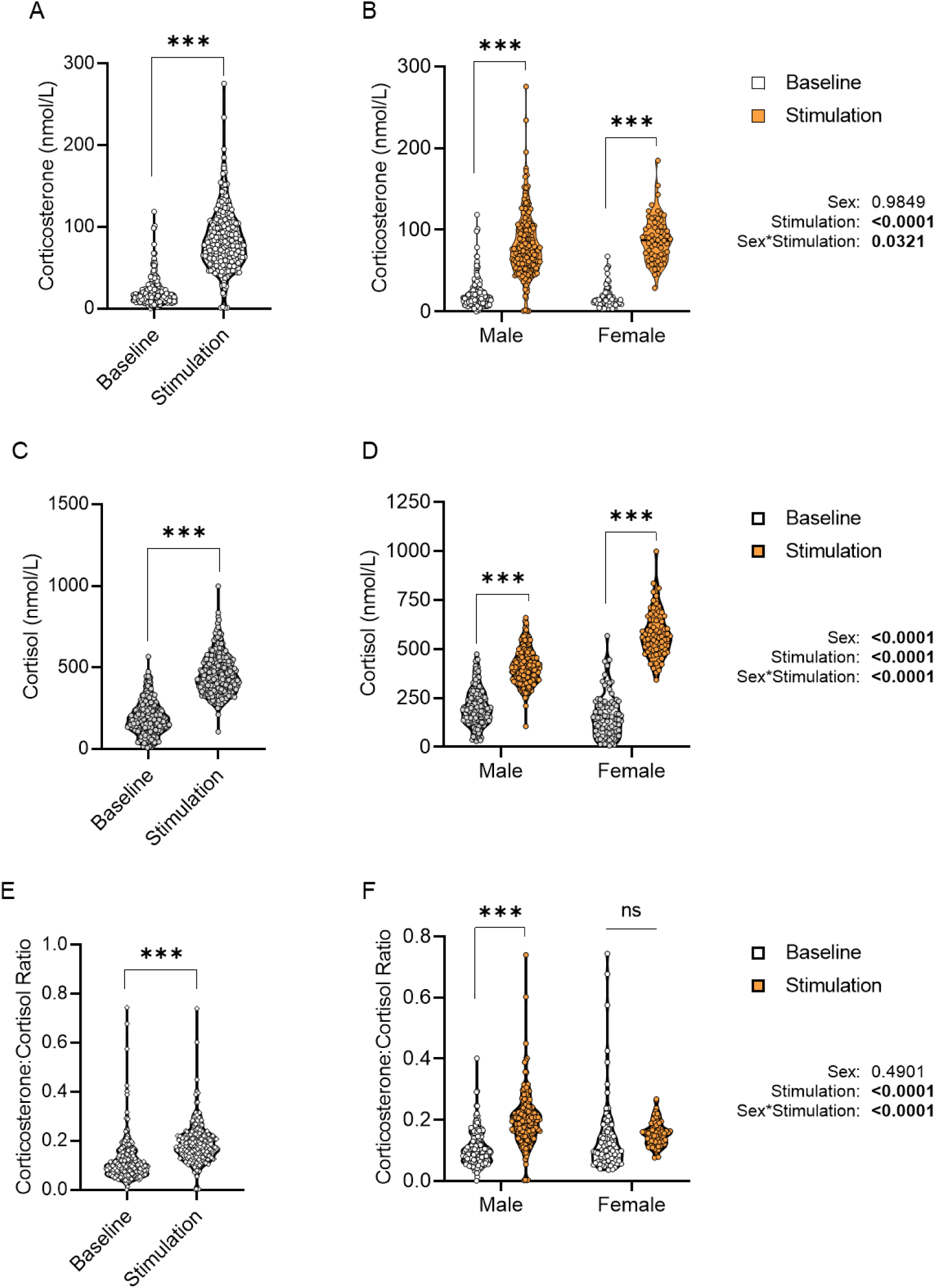
Response to HPA axis suppression and stimulation. Circulating levels of glucocorticoids in healthy subjects following overnight dexamethasone-suppression (“Baseline”) and 30 min after stimulation with ACTH_1-24_ (“Stimulation”). A, Plasma corticosterone. B, Plasma cortisol. C, Corticosterone: cortisol ratio in plasma. Combined data is shown in A, C, E. Data presented by Sex in B, D, F. Data are Mean ± SEM. N=279. Data analysed by paired t-test (A,C,E). Data analysed by Mixed-effects model (B,D,F). \*\*\**P*<0.001

Notably, this stress-induced shift in glucocorticoid balance proved to be sexually dimorphic. When stratified by sex, the increase in the corticosterone:cortisol ratio following ACTH_1-24_ stimulation was observed exclusively in male subjects; this relative shift was not significant in females (Figure 5F). While the relative increase in corticosterone following stimulation was higher in females than males (Figure 5B), females also exhibited a greater absolute increase in cortisol in response to HPA axis stimulation compared to males (Figure 5D), meaning the corticosterone:cortisol ratio remained similar.

## Discussion

The successful translation of corticosterone into a metabolically safer glucocorticoid replacement therapy requires the establishment of a definitive pharmacokinetic framework. We recently demonstrated the clinical promise of this approach in patients with CAH, where acute corticosterone infusion achieved similar suppression of the HPA axis and adrenal androgens without inducing the acute hyperinsulinemia or impaired glucose clearance observed with hydrocortisone, demonstrating proof of concept that matched efficacy may result in reduced toxicity ^30^. However, advancing from this acute proof-of-concept to a viable chronic therapy requires a fundamental understanding of its basal physiological dynamics and absolute elimination kinetics. The present studies establish this essential framework. Our findings highlight that while corticosterone shares many similarities with cortisol regarding its diurnal rhythm and the hepatic enzymes governing its clearance, it is characterised by a faster *in vivo* turnover. This rapid clearance affords a highly dynamic circulating profile with more rapid fluctuations, proving particularly sensitive to changes in HPA axis activation. Together, these data provide the key pharmacokinetic parameters necessary to design appropriate therapeutic regimens and guide future formulation development.

A central finding of this study is the elucidation of the distinct enzymatic pathways driving corticosterone’s rapid turnover. The principal route of clearance of glucocorticoids is hepatic metabolism. Here, we establish that the A-ring of corticosterone is more susceptible to enzymatic reduction than cortisol. *In vitro* metabolism by human hepatic 5β-reductase/3α-HSD proceeded approximately 4 times more quickly for corticosterone than cortisol, although that of 11βHSD2 did not differ. Although the *in vitro* assay is unable to measure 5α-reduction, our *in vivo* urinary metabolite analysis revealed that corticosterone metabolism by 5α-reductase substantially exceeds that by 5β-reductase, while cortisol metabolism was similarly proportioned between both enzymes. These data demonstrate that the corticosterone molecule is a highly efficient substrate for both 5α- and 5β-reductase, and provides a mechanistic explanation for the higher clearance rate observed in our pharmacokinetic studies vs cortisol.

Translating these mechanistic findings into future drug formulations requires characterisation of corticosterone’s *in vivo* elimination kinetics. Given the lack of commercially available clinical grade corticosterone, the isotopically labelled tracer D8-corticosterone was used to determine characteristics of distribution and clearance of corticosterone in healthy volunteers, a tracer we have used previously in patients with Addison’s and CAH ^15,30^. Importantly, our *in vitro* analysis was reassuring in that the tracer was not subject to the primary isotope effect in metabolism by 5β-reductase/3α-HSD or 11βHSD2 (Supplementary Figure 2). Clearance of cortisol is quite consistent across previous studies, however estimates for t_½_ and V_d_ can vary considerably, with estimated mean t_½_ between 60 and 130 minutes in healthy volunteers, and V_d_ varying from 20 to 50 L ^31–34^. In healthy volunteers, we show the clearance of D8 corticosterone was 3-5 times faster than cortisol, with a correspondingly rapid elimination t_½_ of 28 minutes. These highly dynamic kinetics have important implications for future clinical development, with the potential requirement for sustained-release formulations of corticosterone to maintain stable circulating concentrations and provide sustained therapeutic exposure.

Beyond basal clearance, the dynamic kinetics of corticosterone position it as rapid responder during the stress response. In our retrospective *in vivo* analysis, pharmacological HPA axis stimulation with ACTH provoked an exaggerated increase in corticosterone relative to cortisol. Notably, this stress-induced shift proved to be sexually dimorphic, both genders demonstrated significant rises in corticosterone but males had a reduced rise in cortisol compared to females ^35^. Interestingly, a number of small studies performed several decades ago in humans across a wide age range found similar results, with a disproportionately higher rise in plasma corticosterone following ACTH stimulation ^8,36–38^, surgical stress ^39^ and insulin-induced hypoglycemia ^38^, but the consequences of this switch have not been actively explored. This type of switch in proportion has also been observed in the animal kingdom. The tuco-tuco rodent switches its cortisol:corticosterone according to the season ^40^, and rabbits appear to switch from corticosterone to cortisol predominance in early life under ACTH influence ^41^. It is not clear why this occurs, but may reflect adrenal dynamics, with saturation of 17-hydroxylase being a potential mechanism in enhancing corticosterone production ^42^. Overall corticosterone may contribute more to peripheral glucocorticoid exposure under acute stress conditions and play a greater role in suppressing the HPA axis in order to return to homeostasis.

We acknowledge there are several limitations in the current study. While our basal diurnal profiling and stable-isotope pharmacokinetic analyses utilised gold-standard LC-MS/MS quantification, our assessment of HPA axis reactivity relied on a retrospective clinical cohort. Consequently, these legacy samples necessitated the use of a validated, specific radioimmunoassay (RIA) for corticosterone quantification, precluding direct methodological parity across all phases of the study. Of note, in most studies using RIA or LC-MS/MS methods of quantification, basal morning corticosterone is between 10-20 nmol/L ^11,43–46^. This is in accordance with our diurnal profiling data (mean 0800 h = 15.8 ± 9.3 nmol/L) quantified by LC-MS/MS. Furthermore, our exploratory assessment of sex differences in basal diurnal rhythms was limited by a small sample size and was not statistically powered to detect subtle chronobiological dimorphism, highlighting the need for larger, dedicated studies in the future.

In conclusion, while corticosterone and cortisol have historically been regarded as functionally redundant or collectively interchangeable, the present data refute this paradigm. Corticosterone possesses a unique, highly dynamic physiological profile characterised by rapid *in vivo* clearance, distinct 5α-reductase-driven hepatic metabolism, and a disproportionate, sexually dimorphic response to acute stress. These unique pharmacokinetic characteristics provide a vital mechanistic framework to explain its observed tissue-sparing effects and superior therapeutic index. Together, the definitive elimination kinetics established here pave the way for the rational design of advanced sustained-release corticosterone formulations, accelerating its clinical development as a safer, targeted replacement therapy for CAH and other glucocorticoid-dependent conditions.

## Funding

This work was supported by a British Heart Foundation Intermediate Basic Science Research Fellowship to MN (FS/18/20/33449), a Wellcome Senior Investigator award to BRW (107049/Z/15/Z) and a Medical Research Council Clinician Scientist fellowship to RHS (MR/K010271/1). For the purpose of open access the author has applied a Creative Commons Attribution (CC BY) licence to any author-accepted manuscript version arising from this submission.

## Acknowledgements

The authors acknowledge the support of the Edinburgh Clinical Research Facility, and the technical expertise of Lynne Ramage and Alice Ostojic.

## Disclosure statement

The authors declare no competing interests.

## Notes

### Competing Interest Statement

The authors have declared no competing interest.

