## Supplementary Material for "Rapid turnover of corticosterone in humans: the role of a second glucocorticoid hormone"

**Supplemental Material**

Supplementary Figures

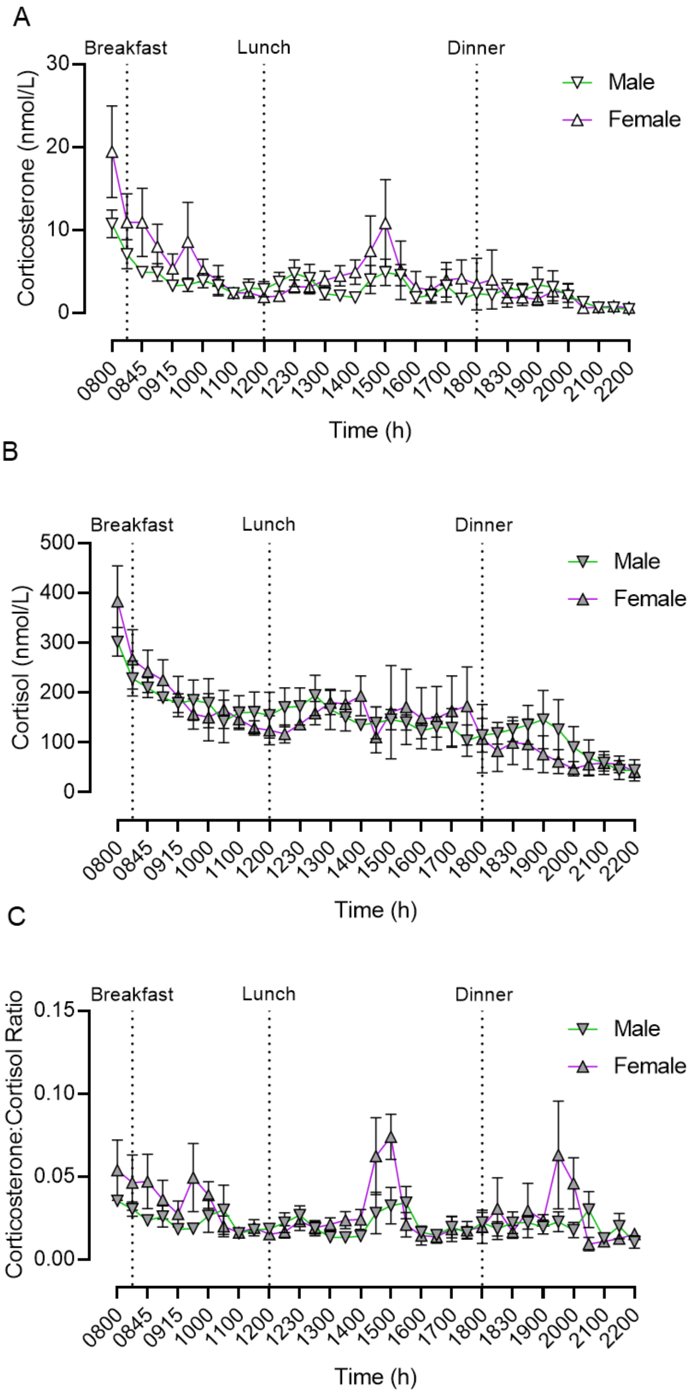


**Supplementary Figure 1.** Diurnal profile of glucocorticoids in healthy male (N=4, green triangle) and female (N=3, purple triangle) subjects. A, Plasma corticosterone. B, Plasma cortisol. C, Corticosterone: cortisol ratio in plasma. Data are Mean ± SEM.

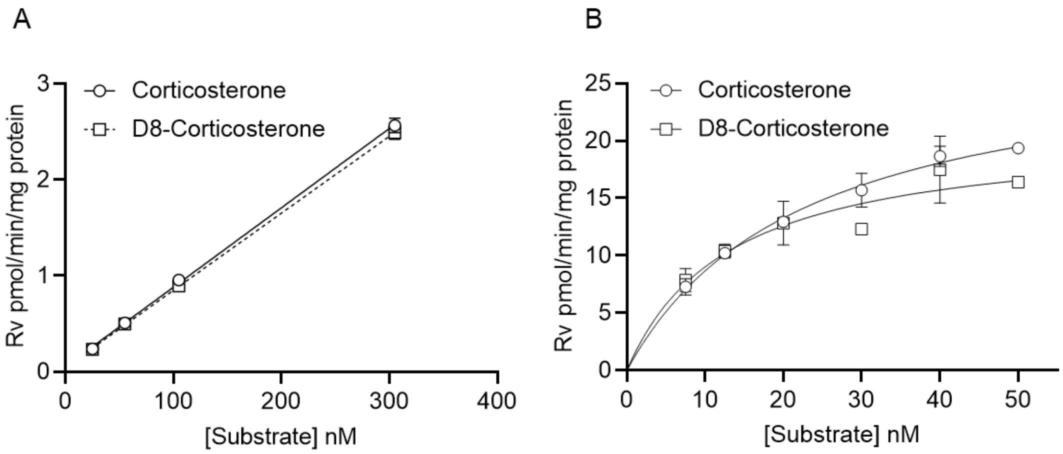


**Supplementary Figure 2**. Metabolism of corticosterone is comparable to D8-corticosterone. A, Metabolism by 5β-reductase/3α-hydroxysteroid dehydrogenase. Reaction velocity (Rv) of corticosterone (open circles) or D8-corticosterone (closed circles) incubated for 120-300 min with human hepatic cytosol (100 µg protein) and enriched with $[^3H]_4$-corticosterone (5 nM). B, Metabolism by 11β-hydroxysteroid dehydrogenase type 2. HEK293-11β-HSD2 cells incubated (30 min) with corticosterone (open circles) or D8-corticosterone (closed circles) and enriched with $[^3H]_4$-corticosterone (2.5 nM). Data are mean ± SEM. N=3.
